# Contrastive Alignment of Expression and Copy Number Highlights Dosage-Insensitive Genes in Cancer

**DOI:** 10.64898/2026.03.01.708901

**Authors:** Garv Goswami, David Xu, Hwi Joo Park

## Abstract

Copy number variations (CNVs) are a hallmark of cancer genomes, yet the relationship between CNV and gene expression is not strictly deterministic. Some genes maintain stable expression despite copy number changes through regulatory compensation. Identifying these dosage-insensitive genes is challenging, requiring methods that distinguish true regulatory escape from technical noise in heterogeneous single-cell data. Here, we present a contrastive learning framework that learns a shared latent space aligning single-cell RNA-seq expression profiles with inferred CNV patterns. Our key innovation is hard negative mining: explicitly training on cell pairs with similar CNV but divergent expression patterns, which represent potential dosage insensitivity. By combining InfoNCE loss with hard negative triplet loss, we learn embeddings where expression-CNV distance quantifies regulatory concordance. We apply this framework to 10 lung adenocarcinoma patients (80k cells) from the GSE131907 atlas, classifying cancer cells as “concordant” (expression follows CNV) or “discordant” (expression escapes CNV). Differential expression analysis between these groups reveals two gene categories: escape genes upregulated in discordant cells despite CNV status, and compensation genes downregulated in discordant cells. Pooled analysis across 40,775 cancer cells identifies significant escape genes including *VSIG4, FCGR1A, TREM2*, and *MARCO*, as well as compensation genes such as *MALAT1, CCL5*, and *CD8A*. These genes represent candidate therapeutic targets and biomarker hypotheses for CNV-independent tumor behavior. Our approach provides a generalizable framework for discovering regulatory escape mechanisms in cancer using standard single-cell RNA-seq data.

## Introduction

Somatic copy-number changes are pervasive in cancer and can alter transcription through gene-dosage effects. However, copy-number alterations do not always translate proportionally into expression modification. Tumors can exhibit transcriptional adaptation and buffering that partially decouples mRNA abundance from the underlying copy number (Bhattacharya et al., 2020). This decoupling motivates the study of CNV–expression “escape” patterns: genes whose expression deviates from dosage expectations at the single-cell level. Such escape behavior may reflect compensatory regulatory mechanisms, cell-state transitions, or selective pressures acting during tumor progression.

Single-cell RNA sequencing (scRNA-seq) enables the interrogation of CNV–expression relationships at cell-level resolution, but quantifying dosage–expression concordance at scale remains challenging in heterogeneous tumors. CNV profiles can be inferred directly from scRNA-seq by aggregating gene expression along genomic coordinates and comparing with reference normal cells, allowing the detection of large-scale chromosomal gains and losses (Tickle et al., 2019). At the same time, gene expression variation is shaped by multiple orthogonal factors, including cell lineage, microenvironmental programs, and dynamic cell states, which complicate efforts to distinguish dosage-consistent from dosage-divergent transcriptional behavior using expression-only representations.

We introduce **Contrastive Learning for CNV–Expression Concordance (CLCC)**, a representation-learning framework that aligns per-cell gene expression profiles with inferred CNV profiles in a shared latent space. The framework is motivated by the observation that dosage-consistent and dosage-divergent transcriptional programs may be difficult to distinguish in expression space alone, particularly in genomically heterogeneous tumors. CLCC draws inspiration from recent successes in cross-modal contrastive learning, most notably CLIP, which learns aligned representations between images and text by pulling together matched modality pairs while separating mismatches (Radford et al., 2021).

In CLCC, a contrastive objective inspired by InfoNCE is used to pull matched expression–CNV pairs together while pushing mismatched pairs apart via negative sampling (van den Oord et al., 2018). To emphasize subtle but biologically meaningful discordance, we incorporate hard negatives and a relative-distance (triplet-style) objective that enforces stronger separation between near-miss mismatches and true matches (Schroff et al., 2015). The resulting embedding geometry induces a simple per-cell *concordance score*, defined by cosine similarity between expression and CNV embeddings, enabling stratification of cells into concordant and discordant regimes and downstream differential expression to identify candidate escape and compensation genes.

We evaluate CLCC on a 10 patient, 80k cell subset of the lung adenocarcinoma (LUAD) single-cell atlas GSE131907 (Kim et al., 2020), which profiles 208,506 cells across 58 samples from 44 patients spanning primary tumors, metastases, and matched normal tissues. Together, CLCC provides a principled pipeline for quantifying CNV-expression concordance across patients and for identifying genes whose regulation systematically departs from expectations held by inferred CNVs.

### Related Works

Multimodal integration has become a central challenge in single-cell analysis. Early methods such as Seurat v3 employed canonical correlation analysis for integrating RNA and ATAC data, but noted that naive concatenation of modalities can be confounded by scale and sparsity differences Stuart et al. (2019). More recent approaches leverage deep learning to learn modality-specific encoders that are aligned in a shared latent space. For example, MultiVI uses a variational autoencoder with separate encoders for RNA and ATAC-seq, enabling coherent embeddings across modalities Lindenbaum et al. (2021). Contrastive learning has emerged as a particularly powerful paradigm for this task: Bian et al. used contrastive alignment of scRNA-seq and scATAC-seq to improve multimodal embeddings Bian et al. (2021), and Lance et al. developed a contrastive framework for RNA and protein modalities that outperformed concatenation and correlation-based baselines Lance et al. (2022).

These works collectively demonstrate that treating each modality with a dedicated encoder, then aligning them in latent space, produces more biologically meaningful and robust embeddings than concatenating raw features. Extending this paradigm to scRNA-seq expression and CNV states offers a natural path forward: per-cell expression vectors and subcluster-level CNV profiles can be projected into a joint representation space, enabling tumor vs. normal separation and the identification of biomarkers driven by both transcriptional and genomic alterations.

### Summary of Findings

#### Cross-modal alignment performance

Our contrastive framework achieved strong alignment between gene expression and inferred CNV modalities. On a held-out test set of 10,000 cells, the model attained a top-1 retrieval accuracy of 87.3% for expression→CNV retrieval and 87.1% for CNV →expression retrieval, with mean reciprocal rank (MRR) values of 0.897 and 0.895, respectively. In both directions, the median rank of the correct match was 1, indicating that matched expression–CNV pairs were typically retrieved as the top candidate. Visualization of the expression– CNV similarity matrix revealed a pronounced diagonal structure, confirming that each expression embedding is most similar to its corresponding CNV embedding and demonstrating successful cross-modal alignment.

#### Discordant cells reveal recurrent escape gene programs

Using the aligned latent space, we classified cancer cells based on the distance between their expression and CNV embeddings, identifying concordant cells whose transcriptional programs closely match CNV expectations and discordant cells whose expression deviates from CNV-driven predictions. Differential expression analysis between discordant and concordant cells revealed a set of *escape genes* that were consistently upregulated in discordant cells in patients, along with *compensation genes* that were down-regulated. Recurrent escape genes included stress-response and immune-related regulators such as *XBP1*, a master regulator of the unfolded protein response. Pooled analysis for all patients further revealed a broad enrichment of myeloid and macrophage-associated genes among escape genes, including *VSIG4, FCGR1A, MARCO*, and *SPI1*, indicating that CNV-expression decoupling is associated with coherent and biologically interpretable transcriptional states.

#### Biological interpretation

These results suggest that discordance between CNV and expression captures a meaningful axis of tumor cell heterogeneity, reflecting transcriptional adaptation, lineage-specific programs, or microenvironmental influences that override copy-number dosage effects. While the limited number of patients limits the overlap of significant genes, the consistency between pooled and patient-stratified results supports the robustness of the identified escape and compensation programs. Importantly, these findings demonstrate that alignment of expression and CNV modalities enables accurate cross-modal retrieval that allow for biologically relevant tasks that would be difficult using either modality alone.

### Dataset and CNV Inference

#### Preprocessing

We obtained raw single-cell RNA-seq UMI count matrices from the lung adenocarcinoma atlas of Kim et al. (2020) (GEO: GSE131907). We selected 10 patients (`P0006, P0008, P0009, P0018, P0019, P0020, P0028, P0030, P0031, P0034`) for which both a tumor sample (`LUNG_T#`) and a matched normal lung sample (`LUNG_N#`) were available.

We focused on these 10 patients as a proof of concept because the availability of a matched normal sample enabled robust CNV inference. In future work, we plan to extend this analysis across additional patients to explore inter-patient variability and validate candidate biomarkers more broadly.

For each patient, cells were filtered to include only those originating from the matched tumor and normal samples. Raw UMI count matrices were loaded and assembled into an AnnData object. We performed basic integrity checks, including verification of unique cell and gene identifiers, confirmation of non-negative integer counts, and assessment of sparsity. Across patients, matrices exhibited the expected high sparsity typical of scRNA-seq data (approximately 94–95% zeros), with no cells containing zero detected genes.

We computed standard per-cell quality control metrics for each patient, including total UMI counts, number of detected genes, and percentage of mitochondrial transcripts. These metrics fell within the expected range for all patients, with no evidence of extreme outliers or technical artifacts. Because of this, we did not apply hard filtering thresholds, retaining all cells to preserve biological heterogeneity relevant to downstream CNV inference and concordance analysis.

Cells were labeled as *Cancer* or *Normal* based on sample origin, with normal lung and normal lymph node samples treated as reference populations for CNV inference. Gene expression profiles and inferred CNV profiles were treated as complementary modalities for subsequent representation learning, without additional cell-type–specific filtering or relabeling.

Finally, the processed AnnData object, including merged annotations and the binary cancer_vs_normal label, was saved for downstream analysis.

#### Copy Number Variation Inference

We inferred copy number variations (CNVs) from single-cell expression data using infercnvpy v0.4.3 icbi-lab (2021–2026), which detects chromosomal amplifications and deletions by comparing tumor cell expression to normal reference cells. Gene expression matrices were first normalized to 10,000 counts per cell and log-transformed. Genes were ordered by chromosomal position using coordinates from GENCODE v44 Frankish et al. (2019), enabling infercnvpy to detect regional CNV patterns through a sliding window approach (window size: 250 genes, step size: 10 genes). Normal lung cells (nLung) served as the diploid reference baseline, with sex chromosomes (chrX, chrY) excluded from analysis.

For each tumor cell, infercnvpy computed a CNV profile representing inferred copy number state across genomic windows. The resulting CNV matrix (cells × genomic windows) contained smoothed output representing copy number relative to diploid.

#### CNV-based Cell Clustering

To identify groups of cells with similar chromosomal alteration patterns, we performed dimensionality reduction and clustering on the CNV profiles rather than gene expression. We applied PCA to the CNV matrix, computed a k-nearest neighbors graph in CNV-space, and performed Leiden clustering (resolution = 0.5) to obtain CNV-defined subclusters. This yielded 20-25 subclusters per patient, each representing cells with distinct chromosomal alteration signatures.

For each CNV subcluster, we computed the mean CNV profile by averaging CNV values across all member cells. These mean profiles serve as reference CNV states for downstream analysis, representing the expected copy number landscape for cells within each subcluster. The CNV sub-cluster assignments and mean profiles were used as anchors in the subsequent contrastive learning framework to identify genes exhibiting expression-CNV discordance.

**Figure 1.**
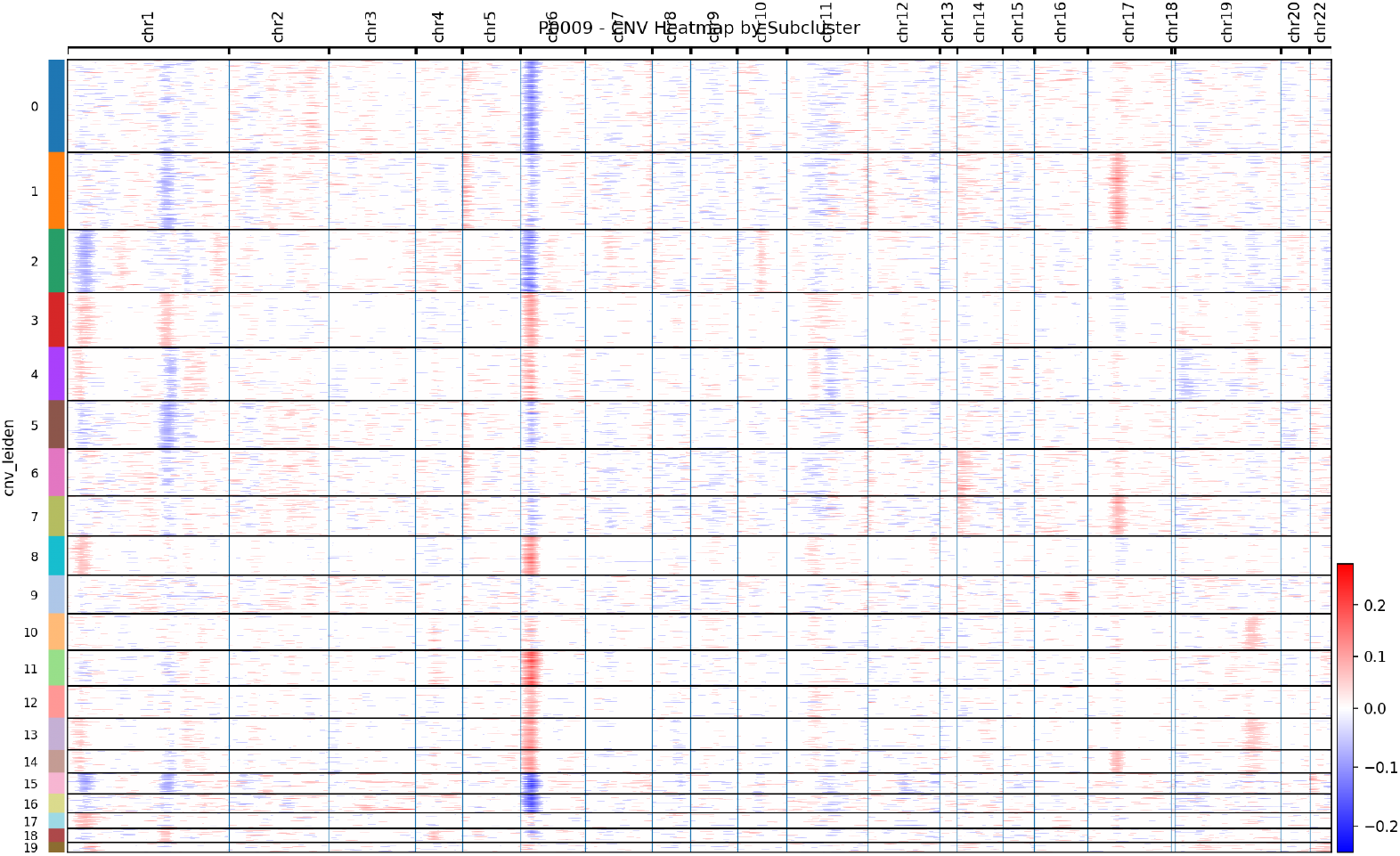
Copy number variation landscape in lung adenocarcinoma. Heatmap showing inferred CNV profiles across the genome for a representative patient. Each row represents a single cell, organized by CNV-defined subclusters (left annotation). Columns represent genomic windows ordered by chromosomal position. Colors indicate inferred copy number relative to diploid: blue = deletion (<2 copies), white = diploid (∼2 copies), red = amplification (>2 copies). Normal reference cells (bottom) show minimal CNV signal, while tumor cells exhibit diverse chromosomal alteration patterns. Distinct CNV subclusters capture cells with similar amplification/deletion signatures.

For our downstream multimodal analysis, we treated the inferred CNV profiles from inferCNVpy as a complementary modality to the single-cell gene expression profiles. Each cell from each patient was thus represented by both its normalized transcriptomic profile and its corresponding inferred CNV signal, together with a subcluster label for evaluation.

#### Dataset Construction

For model training, we implemented a custom PyTorch `Dataset` class (`HardNegativeDataset`) that manages the pairing of expression and CNV data along with precomputed hard negative samples. For each cell *i* in the dataset, the class returns:

- Expression vector: 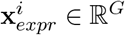 (normalized gene expression profile)
- CNV vector: 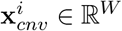 (normalized CNV profile for the cell)
- Hard negative indices: Set of cell indices ℋ_*i*_ with similar CNV but different expression
- Hard negative data: Expression and CNV vectors for cells in ℋ_*i*_

This design ensures consistent pairing of the two modalities at the cell level while enabling efficient access to hard negative samples during training.

**Figure 2.**
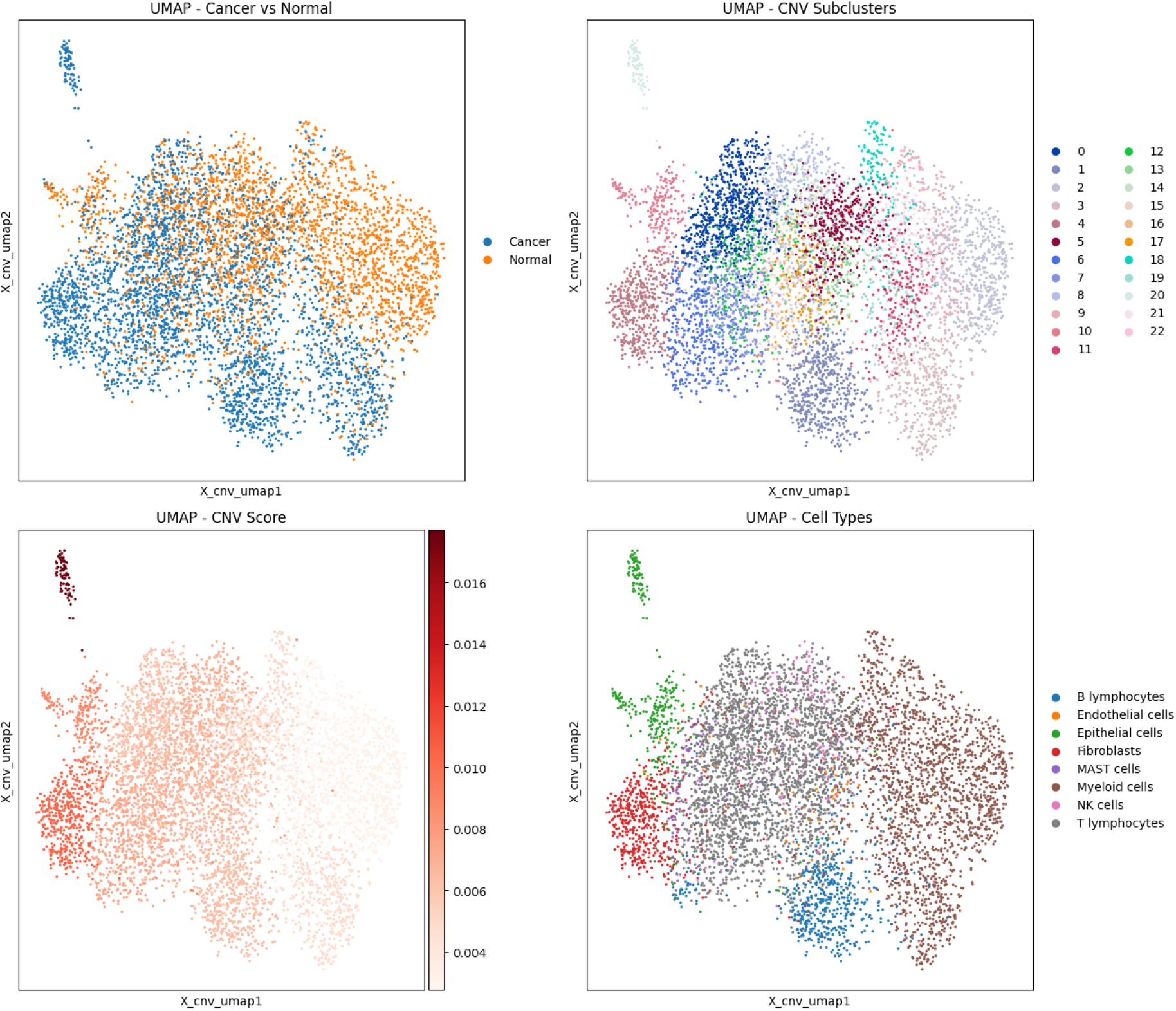
UMAP visualization of inferred CNV profiles across lung adenocarcinoma single cells for patient P0006. (**Top left**) Cancer versus normal cell labels projected onto the CNV embedding, showing partial but incomplete separation. (**Top right**) CNV-based subclusters identified by unsupervised clustering in CNV space, revealing substantial intra-tumoral heterogeneity. (**Bottom left**) Continuous CNV burden score overlaid on the same embedding, illustrating a gradient of aneuploidy rather than discrete CNV states. (**Bottom right**) Annotated cell types projected onto CNV space, demonstrating that CNV structure is largely orthogonal to canonical cell-type identity. Together, these views motivate the need for alignment-based approaches to relate CNV patterns to gene expression beyond cell-type or malignancy labels.

#### Hard Negative Precomputation

Prior to model training, each cell is characterized by a raw expression vector (ℝ^*G*^), raw CNV vector (ℝ^*W*^), CNV subcluster assignment, and cell ID. To identify hard negatives - cells that should exhibit similar expression (due to similar CNV) but do not - we performed pairwise similarity comparisons across all training cells pooled from all patients.

For each cell *i*, we computed cosine similarities between its raw CNV vector and all other cells’ CNV vectors, identifying candidates with CNV similarity > 0.5 as having comparable chromosomal alteration patterns. Among these CNV-similar candidates, we then computed expression similarities and selected those with expression similarity < 0.3 (indicating highly dissimilar expression profiles) as hard negatives. This criterion captures cells where similar chromosomal alterations fail to produce similar expression patterns, suggesting dosage insensitivity or compensatory regulation. For each cell, we identified and cached the top 10 hard negatives ranked by expression dissimilarity. During training, 5 hard negatives were randomly sampled from this pool for each cell in each epoch, providing implicit augmentation through varied hard negative exposure.

To maintain computational efficiency, hard negatives were identified once during dataset initialization rather than recomputed each epoch. For large datasets (> 5, 000 cells), we employed memory-efficient mini-batch sampling, comparing each cell against a random subset of 5,000 candidate cells rather than the full dataset, reducing memory complexity from *O*(*n*^2^) to *O*(*n*·*k*) where *k* = 5, 000. The hard negative indices were cached in a dictionary structure for rapid retrieval during training.

#### Data Loading and Batching

Hard negative mining was performed as preprocessing step prior to training. To control memory usage during pairwise similarity computation, cells were processed in fixed-size anchor batches (typically 200 cells per batch), where each anchor cell was compared against all cells in the dataset to identify hard negatives based on CNV similarity and expression dissimilarity thresholds. This batching strategy was used solely for efficient computation and did not affect the training batch size.

After hard negative precomputation, the dataset was wrapped in a standard PyTorch `DataLoader,` which yielded individual anchor cells along with their precomputed hard negative indices. A custom collation function padded hard negative tensors to uniform size within each batch, as cells may have varying numbers of qualifying hard negatives. A helper function `has_hard_negatives` was used to compute a mask to be used in loss computation to prevent actual computation of zero values. Cells for which no qualifying hard negatives were identified were retained in the dataset and contributed only positive pairs during contrastive training.

#### Data Augmentation

No explicit data augmentation was applied to gene expression or CNV profiles. Unlike image or signal data, individual dimensions of these high-dimensional biological vectors correspond to specific genes or genomic regions, making standard augmentation techniques (e.g., random cropping, rotation, or noise injection) biologically inappropriate. Regularization was instead provided implicitly through contrastive training with hard negatives, which encouraged robust representations without altering the underlying biological measurements.

## Methods

### Model Architecture

#### Encoder Architectures

The framework consists of two parallel encoder networks mapping high-dimensional input data to a shared 128-dimensional latent space.

#### Expression Encoder

Gene expression profiles (ℝ^*G*^ where *G*≈30, 000) are encoded via a feed-forward network: Linear(*G* → 512) → BatchNorm → ReLU → Dropout(0.1) → Linear(512 → 256) → BatchNorm → ReLU → Dropout(0.1) → Linear(256 → 128). The output is L2-normalized to unit vectors.

#### CNV Encoder

CNV profiles (ℝ^*W*^ where *W*≈1, 500 genomic windows) are encoded via a shallower network: Linear(*W* → 256) → BatchNorm → ReLU → Dropout(0.1) → Linear(256 → 128) → BatchNorm → ReLU →Dropout(0.1) → Linear(128 → 128), followed by L2-normalization. The reduced depth reflects the lower dimensionality and smoother structure of CNV data compared to expression.

#### Loss Functions

Training our multimodal encoder requires defining an objective that quantifies how well the model aligns expression and CNV representations. Below, we describe the two loss terms used in our framework, beginning with biological intuition and then providing technical detail.

#### Contrastive Loss (InfoNCE)

At a biological level, the contrastive objective encourages the model to align each cell’s gene expression profile with its *own* inferred CNV profile, so that the shared latent representation reflects coherent genotype–phenotype coupling within the same tumor cell. Concretely, the model is rewarded when the expression embedding of a cell is placed close to the CNV embedding computed from the same cell, and penalized when it is placed close to CNV embeddings from other cells.

Formally, given a minibatch of *B* cells, the encoders map expression and CNV inputs to unit-normalized embeddings **e**_*i*_, **c**_*i*_ ∈ℝ^*d*^. We optimize a symmetric InfoNCE loss that treats matched pairs (**e**_*i*_, **c**_*i*_) as positives and all other pairs in the batch as negatives:

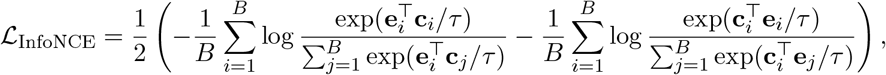

where *τ* is a temperature parameter. This creates a latent space where expression and CNV modalities are aligned at the single-cell level.

#### Hard Negative Mining Loss

While InfoNCE aligns matched expression–CNV pairs, it does not explicitly teach the model what it means for expression to *deviate* from CNV-driven expectations. To emphasize gene-level discordance, we introduce hard negative mining: for each anchor cell *i*, we identify hard negative cells *j* whose CNV profiles are similar to *i* but whose expression profiles are dissimilar (computed in input feature space via cosine similarity after 𝓁_2_ normalization). Biologically, such pairs correspond to cases where two cells share similar underlying CNV patterns, yet exhibit divergent expression programs, suggesting regulatory compensation or escape from dosage effects in at least one of the cells.

Let *j* ∈ ℋ (*i*) denote a mined hard negative for cell *i*, and define cosine distance for unit-normalized embeddings as *d*(**u, v**) = 1 −**u**^⊤^**v**. We apply a margin-based loss that enforces (i) the anchor expression embedding **e**_*i*_ to be closer to its matched CNV embedding **c**_*i*_ than to the hard negative CNV embedding **c**_*j*_, and (ii) symmetrically, the anchor CNV embedding **c**_*i*_ to be closer to its matched expression embedding **e**_*i*_ than to the hard negative expression embedding **e**_*j*_:

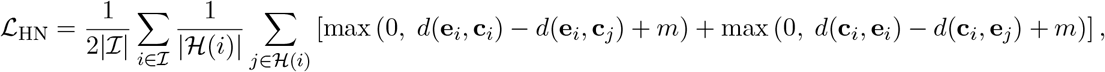

where *m* is a margin hyperparameter and ℐindexes cells for which hard negatives are available. This term explicitly pushes apart pairs that share similar CNV structure but differ in expression, making discordance geometrically meaningful in the embedding space.

**Figure 3.**
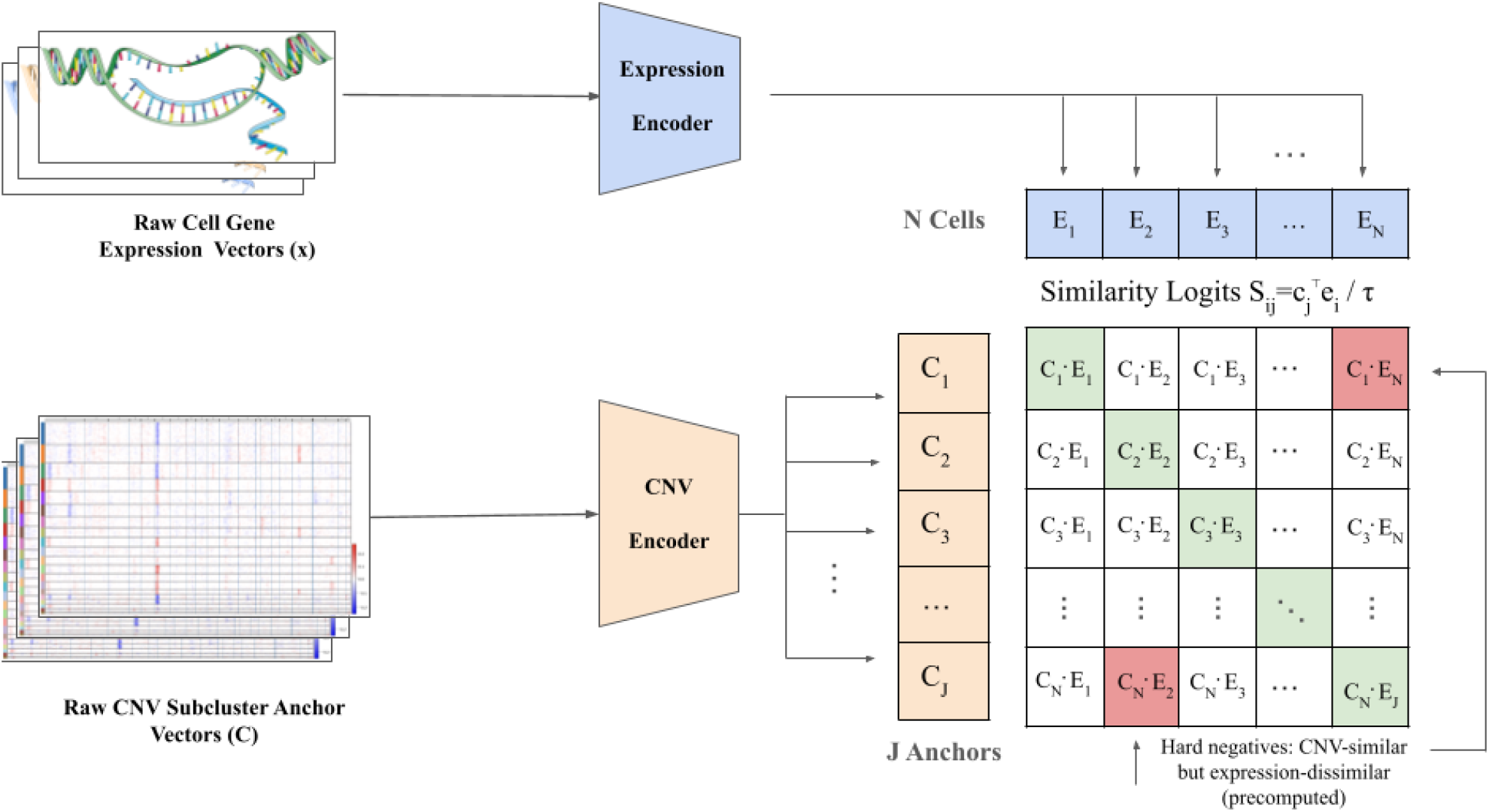
Contrastive pre-training with CNV anchors and hard negatives. Raw scRNA-seq gene expression vectors are encoded to produce per-cell embeddings **e**_*i*_ (top), while CNV subcluster anchor profiles are encoded to produce anchor embeddings **c**_*j*_ (bottom). Similarity logits are computed as 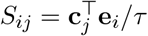and used in an InfoNCE objective, where the positive for cell *i* is its matched CNV anchor *y*_*i*_ (green entries) and other anchors act as negatives. To emphasize expression–CNV discordance, we additionally incorporate precomputed hard negatives (red), defined as cells with high CNV similarity but low expression similarity, and penalize them with a margin-based hard-negative loss.

#### Overall Objective

The final training objective combines standard contrastive alignment with the hard negative penalty:

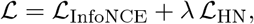

where *λ* controls the contribution of the hard negative mining loss.

### Training Procedure

#### Data Normalization and Splitting

Expression and CNV matrices were z-score normalized across features using training set statistics. Data were split 80%/10%/10% into training (64,462 cells), validation (8,057 cells), and test (8,057 cells) sets.

#### Optimization

Models were trained using AdamW optimizer Loshchilov and Hutter (2017) with learning rate 10^−3^, weight decay 10^−4^, and batch size 256. We employed cosine annealing learning rate scheduling and early stopping (patience: 10 epochs) based on validation loss. Training converged in 80–90 epochs, requiring approximately 30–40 minutes on an Apple M3 Pro GPU using MPS..

#### Hyperparameters

Key hyperparameters were: temperature *τ* = 0.07, hard negative weight *λ* = 0.5, hard negative margin *m* = 0.3, dropout rate 0.1, latent dimension 128.

#### Evaluation Metrics

We evaluated model performance using cross-modal retrieval metrics. For each cell’s expression embedding, we computed its similarity to all CNV embeddings and recorded the rank of the true (matched) CNV embedding. We report the following metrics:

- **Top-***k* **Accuracy**: Fraction of cells for which the correct CNV embedding appears among the top *k* retrieved candidates (*k* ∈ {1, 5, 10}).
- **Mean Reciprocal Rank (MRR)**: The average reciprocal rank (1*/*rank) of the correct match across all cells.
- **Median Rank**: The median rank position of the correct match among all candidates.

We additionally computed an **alignment gap**, defined as the difference between the similarity of the matched expression–CNV pair and the similarity of the hardest negative (most similar non-matched pair). A positive alignment gap indicates successful separation of matched and unmatched pairs.

All metrics were computed bidirectionally (expression → CNV and CNV → expression) and averaged unless otherwise noted.

## Code and Data Availability

### Code Availability

An archive of the codebase is available at https://zenodo.org/record/18465415. The archive contains all scripts required to reproduce data preprocessing, model training, and evaluation. We will maintain this archive with updated code and documentation corresponding to revised preprint versions.

## Data Availability

All data analyzed in this study are publicly available from GEO under accession GSE131907.

## Experiments & Results

### Cross-Modal Retrieval Performance

The contrastive model achieved strong alignment between expression and CNV modalities. On a held-out test set of 10,000 cells, the model achieved a top-1 retrieval accuracy of 87.3% for expression→ CNV retrieval and 87.1% for CNV →expression retrieval, with a mean reciprocal rank (MRR) of 0.896. The median rank of the correct match was 1 in both directions, indicating that matched expression–CNV pairs are typically retrieved as the top candidate.

**Figure 4.**
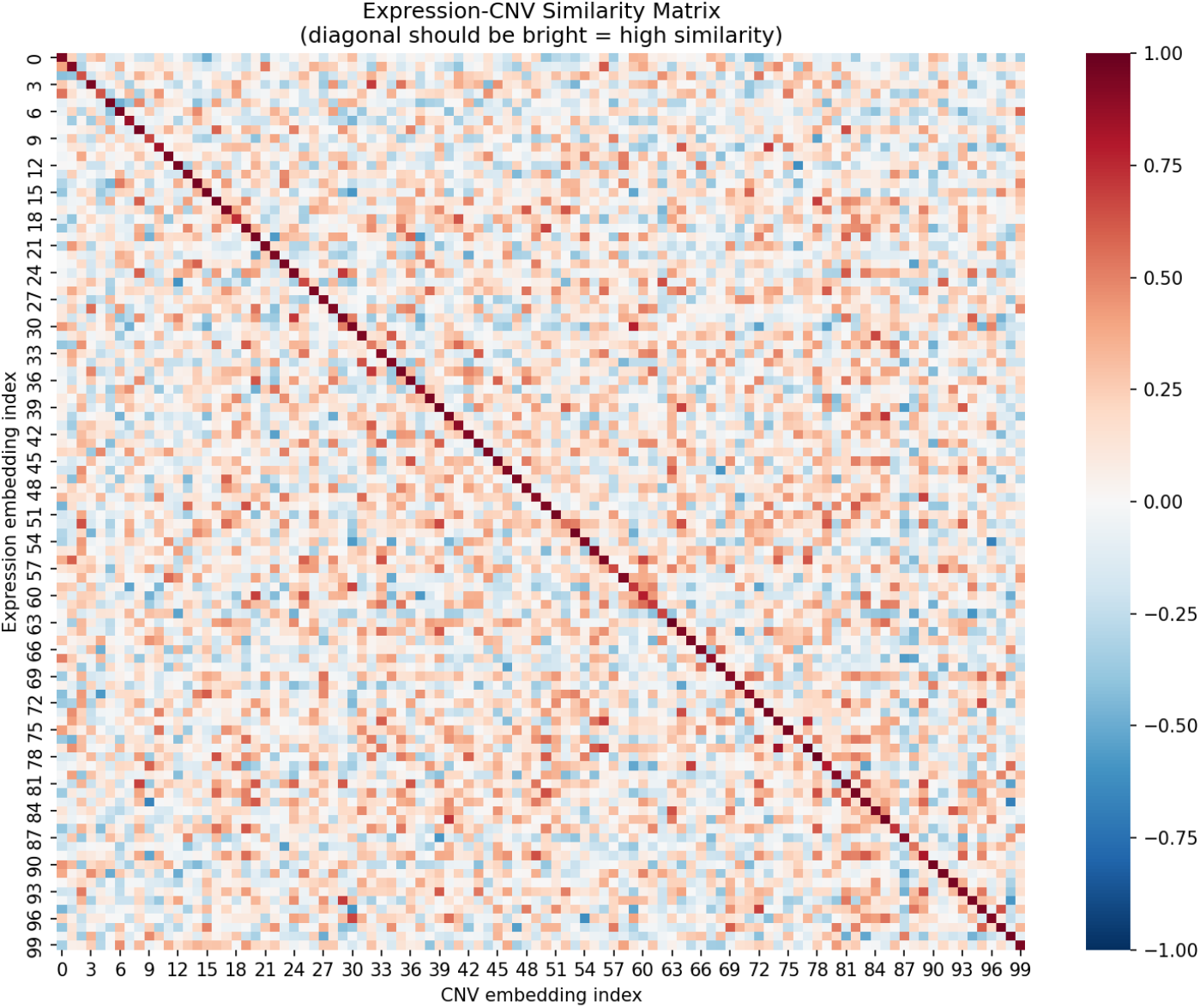
Expression–CNV similarity matrix for a subset of cells. Each entry corresponds to the cosine similarity between an expression embedding and a CNV embedding. The bright diagonal indicates that each expression embedding is most similar to its matched CNV embedding, reflecting successful cross-modal alignment.

To quantify separation between matched and unmatched pairs, we computed the alignment gap, defined as the difference between positive-pair similarity and the hardest negative similarity. The mean alignment gap was 0.108 (expression →CNV) and 0.107 (CNV→ expression), with 87.3% of cells exhibiting a positive gap, confirming reliable discrimination between matched and mismatched pairs.

### Baseline Differential Expression Analyses

To establish context for our contrastive learning-based biomarker discovery, we first performed standard differential expression analyses using established approaches.

### Cancer versus Normal Transcriptional Programs

We compared gene expression between malignant cells (n = 40,775) and normal lung epithelial cells using Wilcoxon rank-sum tests with Benjamini-Hochberg correction. As expected, cancer cells showed significant upregulation of ribosomal protein genes (*RPL41, RPL36, RPLP1, RPS23*) and immunoglobulin genes (*IGKC*), consistent with increased protein synthesis demands in malignant cells. The transcription factor *JUNB*, implicated in cell proliferation and survival, was among the most significantly upregulated genes (log_2_FC = 2.16, FDR ≈ 0).

### CNV Burden-Associated Expression Changes

To identify genes whose expression correlates with overall genomic instability, we stratified cancer cells by total CNV burden (sum of absolute deviations from diploid across all genomic windows). Cancer cells exhibited significantly higher mean CNV scores compared to normal cells (0.0071 vs 0.0053, Mann-Whitney p < 10^−177^), confirming successful CNV inference. Differential expression analysis between high and low CNV burden quartiles revealed widespread upregulation of ribosomal genes and *MALAT1* in high-burden cells, suggesting a global transcriptional response to genomic instability.

### Chromosome-Level Cis-Effects

To validate that inferred CNV profiles correspond to genuine dosage effects, we examined the correlation between chromosome-arm level copy number and mean expression of genes on that arm. Across 21 of 22 autosomes, we observed significant positive cis-correlations (mean Spearman *ρ* = 0.29, range 0.002–0.65). The strongest cis-effects were observed on chromosome 6 (*ρ* = 0.65), chromosome 1 (*ρ* = 0.56), chromosome 11 (*ρ* = 0.44), and chromosome 15 (*ρ* = 0.44). Importantly, cis-correlations consistently exceeded trans-correlations (mean difference = 0.34), confirming that expression changes are driven by local copy number rather than global transcriptional dysregulation.

### Concordance-Based Biomarker Discovery

Having established baseline differential expression patterns, we leveraged our contrastive learning framework to identify genes associated with CNV-expression concordance—a novel axis of tumor heterogeneity invisible to standard approaches.

### Classification of Concordant and Discordant Cells

For each cancer cell, we computed the Euclidean distance between its expression embedding and CNV embedding in the learned joint space. Cells were classified into tertiles: *concordant* (bottom tertile, n = 10,199), *intermediate* (middle tertile), and *discordant* (top tertile, n = 10,199). Concordant cells exhibit gene expression patterns that align with expectations based on their CNV profiles, while discordant cells show expression programs that deviate from CNV-based predictions— potentially reflecting transcriptional adaptation, epigenetic regulation, or microenvironmental influences.

### Per-Patient Concordance Analysis

We performed differential expression analysis between discordant and concordant cells within each of the 10 patients (Table 2). The number of significant genes varied across patients, with P0009 and P0020 showing the strongest concordance signatures (150 and 215 significant genes at FDR < 0.05, respectively). Across all patients, we identified genes consistently classified as escape (upregulated in discordant) or compensation (downregulated in discordant).

**Table 1:** Cross-modal retrieval performance on the held-out test set.

| Metric | Expr→CNV | CNV→Expr |
| --- | --- | --- |
| Top-1 Accuracy | 0.873 | 0.871 |
| Top-5 Accuracy | 0.922 | 0.922 |
| Top-10 Accuracy | 0.941 | 0.942 |
| MRR | 0.897 | 0.895 |
| Median Rank | 1 | 1 |
| Mean Rank | 13.9 | 13.4 |
| Alignment Gap | 0.108 |  |
| Positive Gap Fraction | 87.3% |  |

**Table 2:** Per-patient concordance analysis summary. For each patient, we show the number of cancer cells, concordant and discordant cells (bottom and top tertiles of embedding distance), and significant concordance-associated genes.

| Patient | Cancer | Concordant | Discordant | Sig. Genes | Escape | Comp. |
| --- | --- | --- | --- | --- | --- | --- |
| P0006 | 3,426 | 857 | 857 | 11 | 11 | 50 |
| P0008 | 3,765 | 942 | 942 | 19 | 15 | 4 |
| P0009 | 3,840 | 960 | 960 | 150 | 50 | 18 |
| P0018 | 3,705 | 927 | 927 | 16 | 16 | 50 |
| P0019 | 4,361 | 1,091 | 1,091 | 13 | 12 | 1 |
| P0020 | 4,021 | 1,006 | 1,006 | 215 | 50 | 12 |
| P0028 | 4,705 | 1,177 | 1,177 | 111 | 50 | 3 |
| P0030 | 4,094 | 1,024 | 1,024 | 14 | 14 | 50 |
| P0031 | 5,578 | 1,395 | 1,395 | 12 | 3 | 9 |
| P0034 | 3,280 | 820 | 820 | 49 | 49 | 50 |
| **Total** | **40,775** | **10,199** | **10,199** | — | — | — |

### Recurrent Concordance Genes

To assess the generalizability of concordance-associated genes, we identified genes detected in multiple patients. A total of 22 escape genes and 13 compensation genes were identified in ≥2 patients. The most recurrent escape genes included *CTSB* (cathepsin B, detected in 3 patients, mean log_2_FC = 0.44) and *XBP1* (X-box binding protein 1, detected in 3 patients, mean log_2_FC = 0.38). Notably, *XBP1* is a master regulator of the unfolded protein response, suggesting that discordant cells may activate stress response pathways to cope with CNV-expression mismatch.

Recurrent compensation genes included *CD3D* and *CXCR4* (both detected in 2 patients), suggesting immune-related programs are suppressed in discordant cells despite potential CNV-driven upregulation.

### Pooled Cross-Patient Validation

To maximize statistical power and identify definitive concordance-associated genes, we performed differential expression analysis on pooled data across all patients.

### Pooled Wilcoxon Analysis

Combining all cancer cells (n = 40,775) and stratifying by concordance status (10,199 concordant vs 10,199 discordant), we performed Wilcoxon rank-sum tests on each of 18,965 genes. This analysis identified **5**,**510 genes** significantly associated with concordance at FDR < 0.05.

Applying additional fold-change thresholds, we identified:

- **980 escape genes** (FDR < 0.05, log_2_FC > 0.3)
- **480 escape genes** (FDR < 0.05, log_2_FC > 0.5)
- **467 compensation genes** (FDR < 0.05, log_2_FC < −0.3)
- **226 compensation genes** (FDR < 0.05, log_2_FC < −0.5)

### Top Escape Genes

The most significantly upregulated genes in discordant cells (Table 3) were dominated by macrophage and myeloid markers, including *VSIG4* (V-set immunoglobulin-domain-containing 4; log_2_FC = 1.08, FDR = 7.7 ×10^−61^), *FCGR1A* (Fc gamma receptor Ia; log_2_FC = 1.00, FDR =3.6×10^−56^), and *MARCO* (macrophage receptor with collagenous structure; log_2_FC = 1.09, FDR= 1.8 ×10^−49^). Additional top escape genes included complement components (*C1QC, C1QA*), scavenger receptors (*MSR1, MRC1*), and the transcription factor *SPI1* (PU.1), a master regulator of myeloid differentiation.

**Table 3:** Top 10 escape genes from pooled analysis. Genes significantly upregulated in discordant cells, ranked by statistical significance.

| Gene | Function | $\log_2\text{FC}$ | FDR | % Expressed |
| --- | --- | --- | --- | --- |
| *VSIG4* | Macrophage complement receptor | 1.08 | $7.7 \times 10^{-61}$ | 16.3% |
| *IGSF6* | Immunoglobulin superfamily | 0.90 | $3.1 \times 10^{-58}$ | 18.0% |
| *FCGR1A* | Fc gamma receptor | 1.00 | $3.6 \times 10^{-56}$ | 13.9% |
| *SLC7A7* | Amino acid transporter | 0.84 | $3.4 \times 10^{-54}$ | 17.6% |
| *NCF2* | NADPH oxidase component | 0.89 | $1.3 \times 10^{-53}$ | 18.1% |
| *SPI1* | Myeloid transcription factor | 0.78 | $1.3 \times 10^{-53}$ | 24.4% |
| *HCK* | Src-family kinase | 0.83 | $9.2 \times 10^{-53}$ | 15.6% |
| *CYBB* | NADPH oxidase subunit | 0.79 | $1.4 \times 10^{-52}$ | 19.0% |
| *C1QC* | Complement C1q subunit | 0.89 | $8.1 \times 10^{-52}$ | 23.4% |
| *TREM2* | Triggering receptor on myeloid cells | 0.93 | $4.1 \times 10^{-50}$ | 16.4% |

#### Top Compensation Genes

Genes significantly downregulated in discordant cells (Table 4) included the long non-coding RNA*MALAT1* (metastasis-associated lung adenocarcinoma transcript 1; log_2_FC = −0.19, FDR = −4.3 ×10^−41^), ribosomal proteins (*RPS27, RPL37, RPL30*), and notably, T-cell markers including *CCL5* (log_2_FC = −0.43, FDR = 3.0×10^−10^), *CD8A* (log_2_FC = −0.47, FDR = 1.0×10^−9^), and cytotoxic effectors *GZMH* and *GZMA*.

**Table 4:** Top 10 compensation genes from pooled analysis. Genes significantly downregulated in discordant cells.

| Gene | Function | log<sub>2</sub>FC | FDR | % Expressed |
| --- | --- | --- | --- | --- |
| *MALAT1* | lncRNA, metastasis-associated | -0.19 | $4.3 \times 10^{-41}$ | 99.2% |
| *RPS27* | Ribosomal protein | -0.12 | $6.1 \times 10^{-16}$ | 99.7% |
| *PTMA* | Prothymosin alpha | -0.10 | $2.6 \times 10^{-15}$ | 99.2% |
| *BTG1* | B-cell translocation gene | -0.17 | $2.5 \times 10^{-11}$ | 91.4% |
| *CCL5* | T-cell chemokine (RANTES) | -0.43 | $3.0 \times 10^{-10}$ | 37.2% |
| *CD8A* | T-cell co-receptor | -0.47 | $1.0 \times 10^{-9}$ | 12.0% |
| *GZMH* | Granzyme H | -0.42 | $3.7 \times 10^{-9}$ | 13.6% |
| *GZMA* | Granzyme A | -0.37 | $4.0 \times 10^{-9}$ | 25.7% |
| *NKG7* | NK cell granule protein | -0.41 | $3.5 \times 10^{-8}$ | 23.4% |
| *CST7* | Cystatin F | -0.32 | $7.6 \times 10^{-8}$ | 29.3% |

#### Patient-Stratified Analysis

To account for patient-level batch effects while leveraging the full dataset, we employed a patient-stratified paired analysis. For each gene, we computed the mean expression difference between discordant and concordant cells within each patient, then assessed significance using paired t-tests across the 10 patients. This approach treats each patient as a biological replicate, providing robust inference even when individual patient sample sizes vary.

The patient-stratified analysis identified 3,514 genes with nominal p < 0.05. Top genes by stratified significance included *CD82* (p = 4.9 ×10^−5^), *NRROS* (p = 5.6 ×10^−5^), and *DPP7* (p= 9.5×10^−5^). The concordance between pooled Wilcoxon and patient-stratified results provides additional confidence in the robustness of our findings.

#### Biological Interpretation

The identification of escape and compensation genes provides mechanistic insight into how tumor cells navigate the constraints imposed by their genomic alterations. The striking enrichment of myeloid/macrophage markers among escape genes (*VSIG4, FCGR1A, SPI1, MARCO, TREM2*) suggests that discordant cells—those whose expression deviates from CNV expectations—may be tumor-associated macrophages or cancer cells with activated immune evasion programs.

Conversely, the downregulation of T-cell markers (*CD8A, CCL5, GZMA, GZMH, NKG7*) in discordant cells suggests reduced cytotoxic T-cell infiltration or activity in regions of CNV-expression mismatch. This pattern is consistent with an immunosuppressive microenvironment where genomic instability leads to immune escape rather than immune activation.

Together, these concordance-based biomarkers define a novel axis of tumor heterogeneity that complements traditional mutation-based and expression-based classifications. Cells that successfully escape CNV-expression coupling may represent more aggressive or treatment-resistant sub-populations, with implications for patient stratification and therapeutic targeting.

## Discussion

CLCC identifies transcriptional programs whose expression diverges from local copy-number dosage, revealing biologically coherent patterns of CNV–expression discordance. In lung adenocarcinoma, escape genes were enriched for macrophage and myeloid immune programs, whereas compensation genes were enriched for cytotoxic T and NK cell markers, suggesting that CNV-independent immune regulatory states contribute substantially to transcriptional variation in the tumor microenvironment.

Importantly, CLCC is designed as a representation-learning framework for prioritizing candidate genes and cell states exhibiting dosage-insensitive expression, rather than as a mechanistic model of gene regulation. While the identified escape and compensation genes are supported by strong statistical evidence and biological coherence, their precise functional roles and causal relationships to tumor progression or immune modulation will require orthogonal validation.

Future work may integrate CLCC with perturbation-based assays, spatial transcriptomics, or cross-cancer analyses to dissect the regulatory mechanisms underlying dosage compensation and transcriptional escape. By enabling systematic discovery of CNV–expression discordance at single-cell resolution, CLCC provides a foundation for hypothesis generation and targeted experimental follow-up.

### Limitations

CNV profiles in this study are inferred from scRNA-seq expression, which may introduce shared-signal bias between modalities. Accordingly, CLCC is intended to prioritize candidate dosage-insensitive programs and genes, rather than establish causal CNV–expression relationships. Future work will validate findings on datasets with independently measured CNVs (e.g., paired scDNA+scRNA or orthogonal copy-number assays) and expand benchmarking with additional baselines and targeted ablations of hard-negative mining and loss components.

### Planned updates

In subsequent revisions, we plan to (i) evaluate CLCC on paired multi-omics datasets with independently measured CNVs, (ii) add ablations isolating the contribution of hard-negative mining and the margin-based loss, and (iii) broaden baseline comparisons to include simple dosage-residual models and alternative multimodal alignment objectives.

## AI Use Disclosure

Large language models, including ChatGPT (OpenAI), Claude (Anthropic), and Gemini (Google DeepMind), were used during the development of this work. These tools assisted with brainstorming and conceptualization, clarifying technical ideas, drafting and editing text, and generating illustrative code snippets for data processing and analysis. All model-generated content was critically reviewed, validated, and, where necessary, modified by the authors. The authors take full responsibility for the accuracy, originality, and integrity of the final manuscript.

